# Reanalysis of sequences of alleged Javan tiger highlights the difficulties in studying big cats and the need for high throughput sequencing

**DOI:** 10.1101/2024.04.15.589466

**Authors:** Anubhab Khan, Yulianto Yulianto, Sabhrina Gita Aninta, Wirdateti Wirdateti

**Affiliations:** Section for Computational and RNA Biology, Department of Biology, University of Copenhagen, Denmark; School of Biodiversity, One Health and Veterinary Medicine, University of Glasgow, UK; Department of Biology, Pwani University, Kenya; Research Center for Applied Zoology, National Research and Innovation Agency (BRIN), Cibinong, West Java, Indonesia; Research Center for Biosystematics and Evolution, National Research and Innovation Agency (BRIN), Cibinong, West Java, Indonesia

## Abstract

Big cats are of conservation concern throughout their range. Genetic tools are often employed to study them for various purposes. However, there are several difficulties in using genetic tools for big cat conservation which may be resolved by modern methods of DNA sequencing. Recent reports of discovery of Javan tigers in West Java, Indonesia highlights few of the difficulties in big cat genetics. We reanalysed the data of the original reports and found that the results were unreliable. However, resequencing of the DNA extracts confirm that the sighting could have been that of a tiger, but the subspecies cannot be confirmed. The work highlights the urgency for development of high throughput sequencing infrastructure in the tropics and the need for reliable databases for studies of big cats.

## Introduction

Big cats are rare and elusive. While they attract a huge amount of conservation funding and have been studied across their range by various researchers, they remain a mystery. Poaching, livestock depredation, trafficking, ex-situ breeding and dispersal events are a few examples of the importance of identifying big cats properly. Genetics tools are often used for identifying big cats but the classical tools may be inadequate. A recent study by Wirdateti et al. (2024) where the presence of a Javan tiger has been reported from West Java is one such example. Javan tigers were classified as extinct by IUCN in 2008 (Jackson and Nowell 2008) and their presence has not been detected since the 1990s. However, in 2019, a local resident had seen an alleged tiger near a village in West Java and one of the authors of Wirdateti et al. (2024) collected a hair (Wirdateti et al. 2024).

To determine the possibility of finding the extinct javan tiger, museum samples of Javan and Sumatran tigers were also collected, and DNA was extracted from the hair strand and the museum samples (Wirdateti et al. 2024). They sequenced cytochrome B (cytB) region and performed comparative phylogenetic analysis with other previously published cytB sequences of tigers and leopards to conclude that the hair belongs to a Javan tiger. Here, we have reanalysed those sequences and repeated a few experiments to highlight the difficulties of studying big cat genetics in the wild and some potential solutions.

## Methods

### Phylogenetic tree reconstruction

We reanalysed the Javan tiger sequences in Wirdateti et al. (2024) by making a phylogenetic tree including additional cytB and NuMt sequences downloaded from NCBI (see Supplementary Table 1). The sequences were aligned using MAFFT (Katoh et al. 2002). Three batches of analysis were done based on the length of sequences analysed and the type of sequences used: (1) sequences MH290773, AB211408 to AB211411, and FJ403465 were removed from analysis due to excess of missing data. All regions with any missing data were trimmed using Jalview (Waterhouse et al. 2009). This retained 36 sequences with 265 bp data for analysis. (2) Sequences AB211408 to AB211411, FJ895266, FJ403466, FJ403467, MH290773, and FJ403465 were removed from analysis due to excess of missing data and all regions with any missing data were trimmed using Jalview (Waterhouse et al. 2009). This retained 33 sequences with 971 bp data for analysis. (3) cytB NuMt sequences were aligned using MAFFT and sites with missing data were trimmed using Jalview retaining 453 sites with 8 sequences.

Neighbour joining trees were built using MAFFT online service (Katoh et al, 2017). Default option was chosen for multiple sequence alignment. For building the tree, the option of Conserved sites was chosen which retained 252 sites for dataset (1), 937 sites for dataset (2), and 240 sites for dataset (3). Raw differences were used for Substitution model and bootstrap resampling was performed 1000 times.

### DNA extract re-sequencing

The NuMt sequences got amplified due to stochastic binding of the primers to the NuMt region instead of mitochondrial cytB in Wirdateti et al. (2024); we expect this can be rectified by repeated PCR and sequencing. Some amount of the extracted DNA from the hair strand and the Javan tiger specimen from Museum Zoologicum Bogoriense (Wirdateti et al. 2024) remained in the tubes. We re-amplified and re-sequenced these samples several times as described in Wirdateti et al. (2024). One of the sequencing batches yielded an approximately 900 bp long DNA fragment from the test hair strand and the museum sample. These new sequences were aligned to the sequences listed in Supplementary table 1 using MAFFT and trimmed using Jalview. Two batches of analysis were done due to the trimming. One that retained 38 sequences and 264 bases and the other retained 35 sequences and 907 sites.

## Results

We reanalysed the sequences generated by Wirdateti et al (2024) alongside a nuclear copy of cytB pseudogene (NuMt) sequence of Bengal tiger (Panthera tigris tigris, AF053053.1) and the cytB sequences of several other tiger subspecies. The clustering of the putative Javan tiger sequence (“Test sample” in Figure 1) and the museum sample of Javan tiger (“OQ601562” Figure 1) with the NuMt sequence revealed that the sequences generated for the samples were NuMts and not cytB region that they were being compared to (Figure 1).

**Figure 1:**
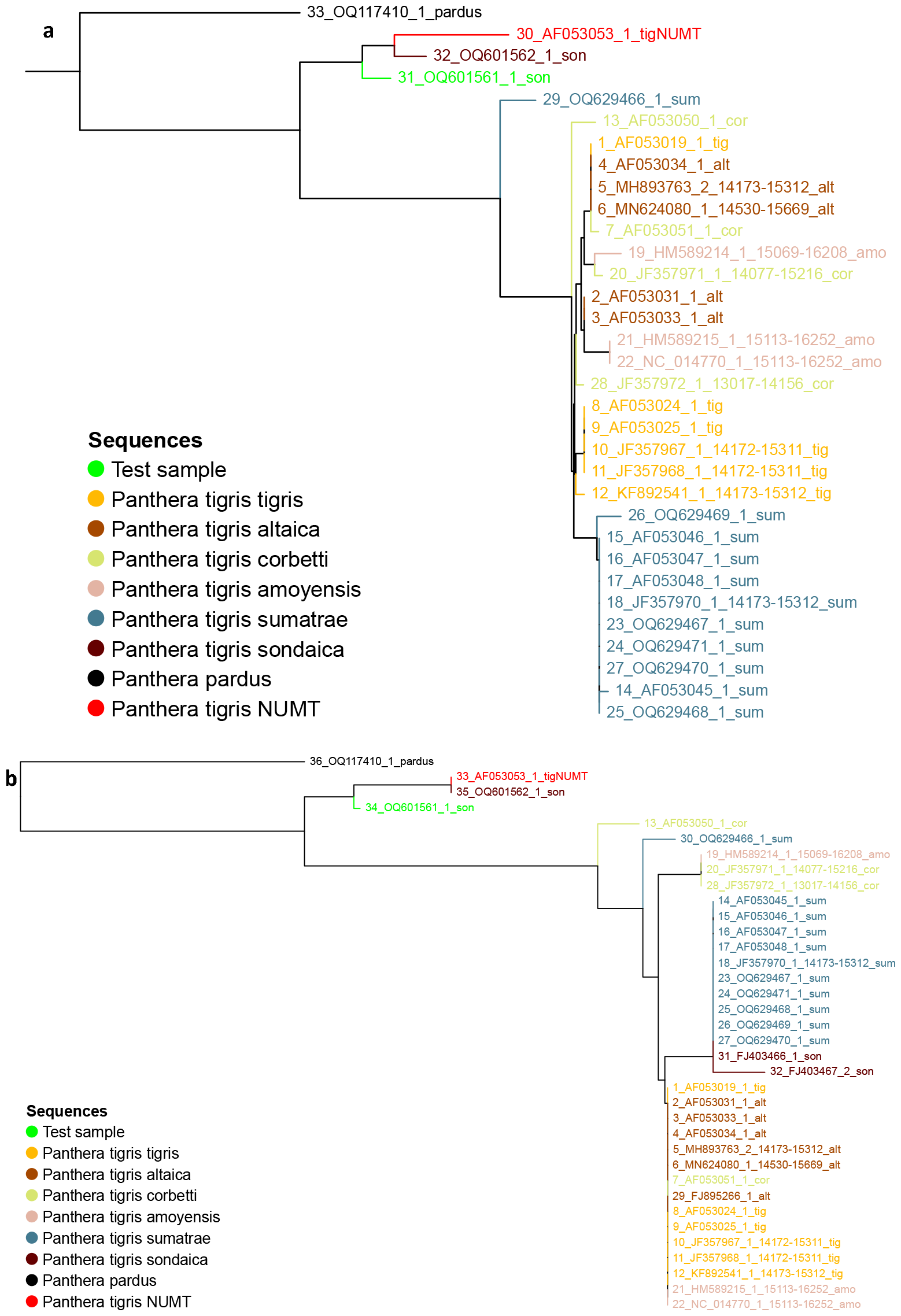
Neighbour joining tree of the test sample (hair strand of putative Javan tiger, green), the javan tiger museum sample (OQ601562_1_son, brown) and other tiger sequences using (a) 36 sequences and 252 sites and (b) 33 sequences and 937 sites. A sequence of nuclear copy of cytB pseudogene (NuMt, red) was included in the analysis.

We further added NuMt sequences of other Pantherine cats and dog (as an outgroup) to see the possibility of NuMts as species identifier (Figure 2). Comparisons of the available annotated cytB NuMt sequences of Pantherine cats on NCBI demonstrates that NuMt sequences can be highly divergent within the Panthera lineage (Figure 2). It is also observed that the Panthera leo cytB NuMt sequence seems more diverged from other cats and from a canid cytB NuMt.

**Figure 2:**
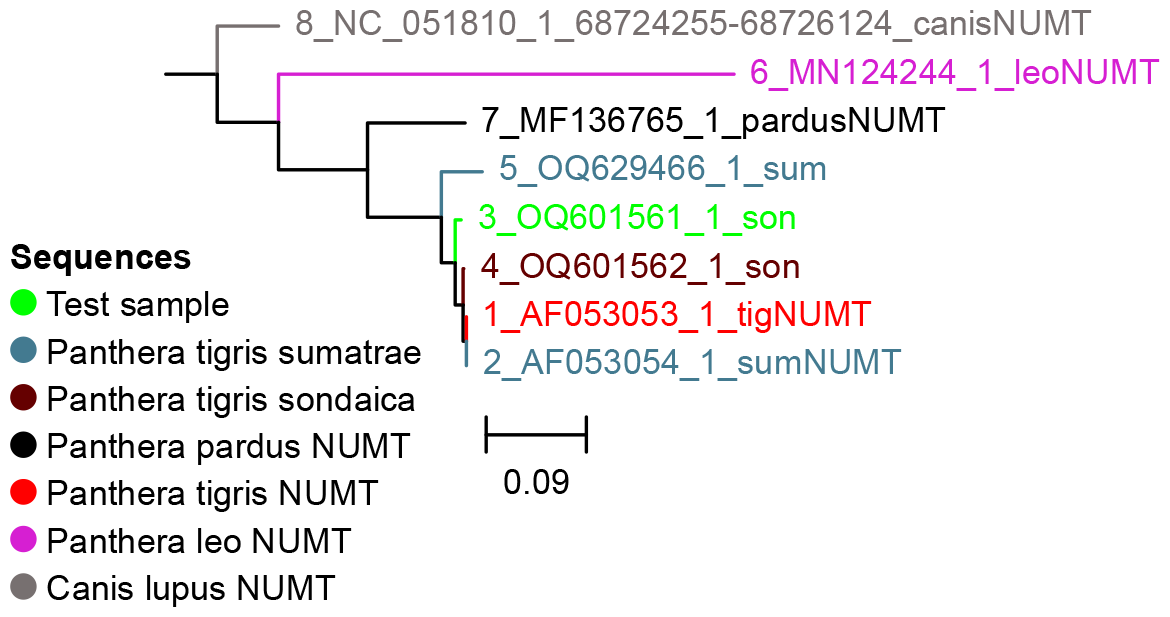
Neighbour joining tree of the test sample (hair strand of putative Javan tiger, green) and cytB NuMt sequences from other big cats and a dog.

We re-sequenced DNA extract remains from the putative Javan tiger hair strand and the specimen of Javan tiger form the museum, and retained sequences closely related to the cytB sequences of the other tigers (Figure 3). However, the polytomy that was made alongside other Sumatran tigers revealed that mitochondrial cytB DNA does not have the power to distinguish between Sumatran tigers and Javan tigers (Figure 3).

**Figure 3:**
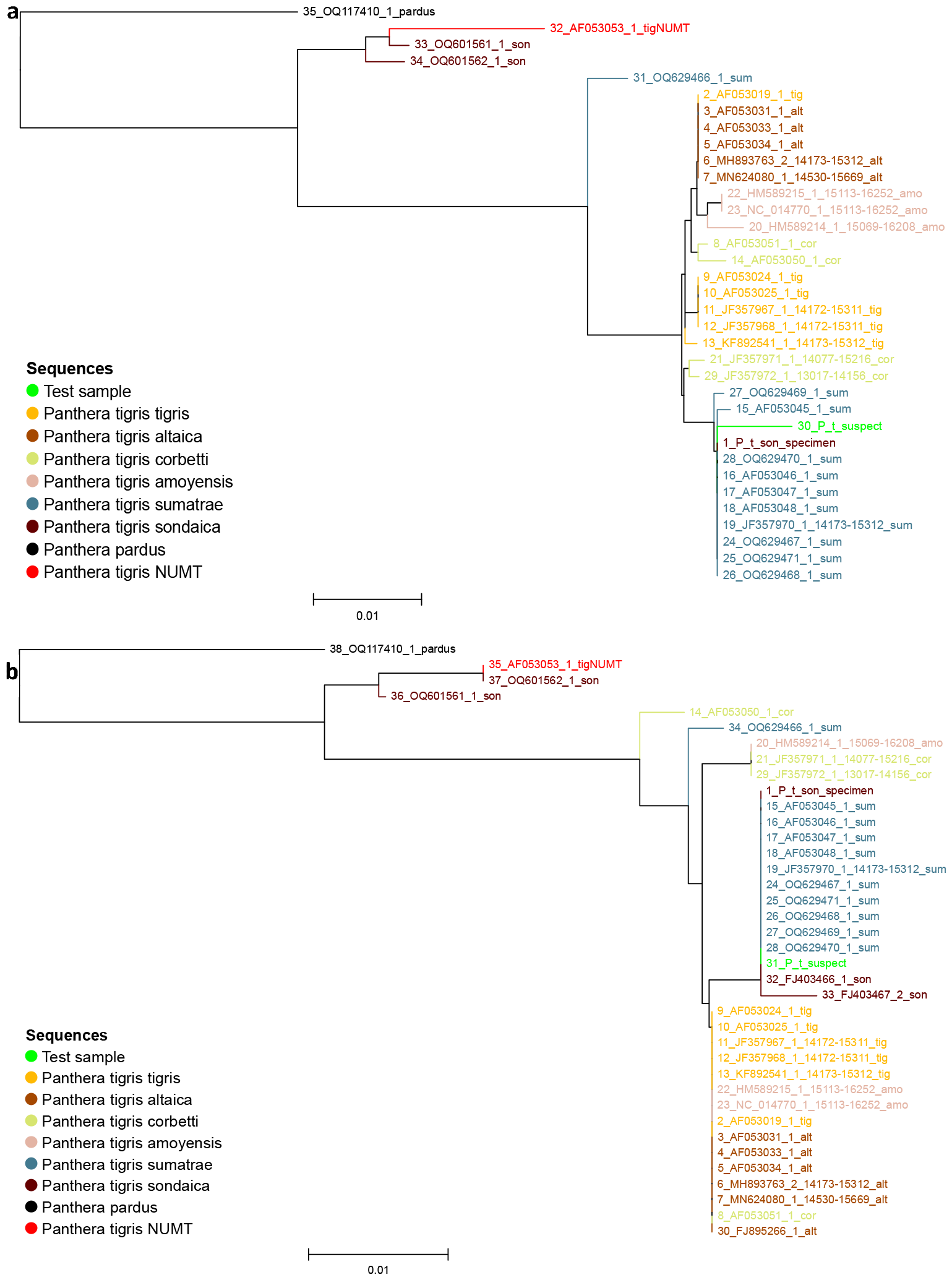
**Neighbour joining tree of the re-sequenced test sample (hair strand of putative javan sample, P_t_suspect, green), museum sample of javan tiger (OQ601562_1_son, brown) and other tiger sequences using (a) 38 sequences and 252 sites and (b) 35 sequences and 869 sites.**

## Discussions

Mitochondrial genomes are used commonly for delimiting species and subspecies. However, they have several limitations especially for big cats. In big cats, nuclear copies of mitochondrial pseudogene have been a common problem for genetic investigations (Kim et al. 2006, Morgan et al. 2021). These regions, often called NuMts, evolve independent of the mitochondrial genome, have different rates of evolution, and should not be compared directly. However, since they have sequence similarities, primers intended to amplify the mitochondrial copy of a gene, may accidently amplify the nuclear copy and provide misleading results.

Additionally, since mitochondrial DNA is inherited matrilineally, analysis of only mitochondrial sequences limits detection of admixture events. For example, mitochondrial DNA would reveal the subspecies of the mother of a tiger sample but says nothing about the father. This is especially relevant if samples of an admixed tiger are obtained for forensic analysis or captive breeding or for tracing the origins of a sample.

Mitochondrial DNA is cheap to obtain and is better preserved in non-invasive samples due to its abundance in the cells compared to nuclear DNA. Since, mitochondrial DNA has been used for a long time, and the protocols are more familiar to most researchers. However, ancestry informative SNP panels (Khan et al. 2022), low depth sequencing (Fuentes-Pardo and Ruzzante 2017), multiplex PCR panels (Natesh et al. 2019) and pooled sequencing (Fuentes-Pardo and Ruzzante 2017) are cheap alternatives that overcome the limitations of mitochondrial DNA from non-invasive samples. Presently, the need for computational infrastructure and expertise, high start-up costs, high import cost of reagents and access to high-throughput sequencers are a major hindrance in the tropical countries (Khan and Tyagi 2021).

Presently, we can confirm that the hair sample observed in West Java nests within the clade of Sundaland tigers but we are unable to assign it to a subspecies. This is partially because of a lack of database of extinct tiger lineages. There are several specimens of Java, Bali, and Caspian tigers in the museum (Yamaguchi et al. 2013), but they are understudied and hence genetic resources from these lineages are lacking. This has limited our ability to explore the possibility of shared haplotypes in extant tigers and to determine the similarities and differences between extinct and extant lineages. Wilting et al. (2015) for example demonstrate the inability of short DNA sequences to resolve the tiger subspecies while whole genome sequencing studies (Liu et al. 2018, Armstrong et al. 2020) have been successful in doing so at least for the extant lineages.

We would like to highlight few important challenges in big cat genetics revealed by the study in Wirdateti et al. (2024):

A. Good quality samples of big cats are difficult to come by and non-invasive samples are the norm. Due to the abundance of mitochondrial DNA compared to nuclear DNA in cells, it has been common practice to analyse mitochondrial sequences. However, NuMts are common in big cats and can lead to misrepresentation in analysis.
B. Despite there being methods for analysing whole genomes from non-invasive samples like shed hair and faeces (Khan et al. 2020, Tyagi et al. 2022, Khan 2023) lack of resources in terms of sequencing facilities, computational infrastructure, DNA enrichment reagents or SNP panel makes it an inaccessible tool (Khan and Tyagi 2021).
C. There is an urgent need to develop expertise in next-generation sequencing in biodiversity rich tropical countries otherwise we will miss out on discovering species. This is especially important now in the on-going mass extinction of the Anthropocene.
D. Big cats and their parts are often the subject of forensic investigation in cases of trafficking, livestock depredation or like in the case of Wirdateti et al (2024). All such investigations could be affected by NuMts if analysis is only restricted to the mitogenome. There is an urgent need for cost effective SNP panels for ancestry determination and population assignment (for example Khan et al. 2022)
E. Databases like GenBank and the journals need to insist and verify that the sequences reported are archived properly and made available. This would make it easier for the peers to build on the science for big cats. For example, despite their being so many genetic studies on tigers, the sequences are difficult to retrieve or use because of improper annotations or not archiving (for example sequences from Wilting et al. (2015) need to better annotations and genome assemblies from Armstrong et al (2021) and Zhang et al. (2023) need to be released and many more).

